# Metagenomics reveals potential interactions between CPR bacteria and their phages in groundwater ecosystems

**DOI:** 10.1101/2025.02.25.640227

**Authors:** Bingxin Hu, Liyun An, Mengdi Wu, Jinbo Xu, Yong Nie, Xiao-Lei Wu

## Abstract

Candidate phyla radiation (CPR) is a vast lineage composed of bacteria with ultra-small size, streamlined genomes, notable defects in core metabolic potential and symbiotic lifestyle, which are widely detected in groundwater ecosystems. Increasing attention has focused on the physiological and ecological significance of CPR bacteria, while the potential interactions between CPR bacteria and their phages still need more exploration. Here, we collected 82 groundwater metagenomic datasets and further derived 1,162 phages with the potential to infect 2,439 groundwater CPR metagenome-assembled genomes (MAGs). Notably, the groundwater CPR MAGs were predominantly infected by temperate phages, and the abundance profiles of phage-CPR interactions in groundwater ecosystems supported the ‘Piggybacking the Winner’ (PtW) hypothesis, suggesting that lysogenic infection was a strategy co-evolved by CPR bacteria and their phages in groundwater ecosystems. Intriguingly, the groundwater CPR phages encoded various auxiliary metabolic genes (AMGs) that might promote symbiotic lifestyle and metabolic potential of host CPR MAGs. These included AMGs associated with concanavalin A-like lectin/glucanases superfamily and O-Antigen nucleotide sugar biosynthesis, which could enhance surface adhesion of host CPR MAGs. Moreover, AMGs related to ABC transport system and P-type transporter could strengthen metabolic exchange and uptake of essential nutrients from surroundings. Additionally, AMGs involved in various metabolic pathways might alleviate metabolic deficiencies in host CPR MAGs. This study elucidated the mechanisms of phage-CPR interactions and the potential roles of phages in modulating the physiology and ecology of CPR bacteria within groundwater ecosystems.

## Introduction

Aquifers are essential components of the Earth’s deep terrestrial subsurface, also serving as significant natural reservoirs. They supply approximately 30% of the world’s freshwater [1], playing vital roles in drinking water provision, agricultural irrigation, and various industrial processes [2]. The important roles of groundwater in human society and the Earth’s environment call for deeper understanding of groundwater ecology and geochemistry [3]. Despite having nutrient-limited oligotrophic conditions, groundwater ecosystems can still provide niches for various microbes (including bacteria and archaea) [4, 5]. Groundwater microbial communities make great contributions to subsurface biogeochemical cycling of key elements (such as carbon, nitrogen, sulfur, phosphorus, and numerous metals) and energy conversion [6].

Notably, groundwater ecosystems with nutrient-limited and anoxic environmental conditions provide ideal habitats for a keystone taxon with unique evolutionary status and unclear ecological functions — CPR bacteria [7–11]. CPR bacteria comprise over 50% of all bacterial diversity [12]. The discovery of CPR bacteria has reshaped our understanding of the tree of life and underscored the critical roles of uncultured bacteria in the environment [13]. Despite their high diversity, CPR bacteria exhibit conserved traits, including ultra-small cell size, streamlined genomes, and minimal biosynthetic capabilities [14–16]. Thus, CPR bacteria adopt lifestyles relying on other cells, either by episymbiotically attaching to larger host bacteria to get nutrients, or by deriving essential compounds from surroundings [16].

Viruses are the most abundant, widespread and genetically diverse biological entities in the biosphere, with a conservative estimate above 10^31^ [17]. Recent studies have indicated that viruses widely inhabit in aquatic [18–20], terrestrial [21], human-associated [22, 23] and engineered environments [24], even in extreme environments such as desert [25], glacier [26], permafrost [27] and acid mine drainage [28]. Viruses profoundly influence the composition and evolution of microbial communities, playing crucial roles in various ecosystems [29, 30]. Virulent viruses can regulate the composition of microbial communities by targeting and lysing the dominant populations, and contribute the host-derived organic matter to surroundings through cell lysis, thereby affecting the global biogeochemical cycle [31]. Moreover, viruses can regulate metabolism and physiology of their host microbes via auxiliary metabolic genes (AMGs) [32]. Virulent viruses exhibit AMGs associated with host metabolism to boost progeny reproduction, while temperate viruses tend to encode AMGs related to host physiological regulation to benefit virus-host coexistence [33]. The ecological roles of virus-host interactions in different ecosystems requires thorough contemplation.

Recently, some studies have focused on phages that can infect CPR bacteria [34, 35]. Due to the increasing relevance to human oral health and disease, Saccharibacteria, Gracilibacteria and Absconditabacteria (SGA), lineages within CPR bacteria, have attracted much attention. Liu et al. explored the diversity of CRISPR-Cas systems in SGA, and sought phages with the potential to infect them by matching their CRISPR spacer inventories to phage databases (IMG/VR v3 and GVD Human Gut Virome).

By examining the genetic code of predicted SGA phages, they found that some of them might infect both standard and alternatively coded host bacteria [34]. Moreover, Wu et al. establish the first metagenomic Groundwater Virome Catalogue (GWVC) and explored the diversity, lifestyle, and functional potential of phages connected to the keystone ultrasmall symbionts (CPR bacteria and DPANN archaea). They found that partial CPR phages harbored genes associated with cell-surface modification that potentially assist symbiont cells adhere to free-living microbes [35]. However, the mechanisms underlying phage-CPR interactions, including the relationships between their relative abundance across different ecosystems and the roles of phages in supporting the symbiotic lifestyle and metabolic potential of CPR bacteria through AMGs, remain underexplored. Here, we analyzed 82 groundwater metagenomic datasets that have been reported to contain abundant CPR bacteria, and further explored the diversity and novelty of CPR phages, as well as the mechanisms underlying phage-CPR interactions in groundwater ecosystems.

## Materials and Methods

### Data acquisition and processing

Based on three previous studies [7, 8, 11], a total of 82 groundwater metagenomic data sets were downloaded from NCBI Sequence Read Archive and NCBI BioSample databases (Data S1). Raw reads were trimmed using the Trimmomatic v0.3962 [36] (parameters: LEADING:2 TRAILING:2 SLIDINGWINDOW:4:20 MINLEN:50), following by individually assembly using SPAdes v3.15.4 [37, 38] (parameters: -meta, -k 21,33,55,77,99,127) or MEGAHIT v1.2.9 [39] (default parameters). Moreover, metagenomic-assembled genomes (MAGs) binned from the 82 groundwater metagenomic data sets were also downloaded. After taxonomic assignment using GTDB-Tk v1.5.0 [40] based on Genome Taxonomy Database (GTDB, http://gtdb.ecogenomic.org), 2,790 CPR MAGs were retained for further analysis.

### Recover of groundwater viral contigs

Viral contigs were recovered from assembled contigs using VirSorter v2.1 [41] (parameters: --exclude-lt2gene), only viral contigs ≥ 5 kb were retained. Completeness of the initial viral contigs was estimated using the CheckV v0.8.1 [42] pipeline (end_to_end program). The final viral contigs were obtained by merging the output file of CheckV (‘viruses.fna’ and ‘proviruses.fna’) which had removed contamination of host microbes. Any viral contigs annotated as ‘no viral genes’ were also manually discarded. Completeness and length of the final viral contigs was also estimated using the CheckV v0.8.1 pipeline (end_to_end program). The groundwater viral contigs were further clustered into virus operational taxonomic units (vOTUs) with parameters of 95% average nucleotide identity (ANI) and 85% alignment fraction based on scripts provided in CheckV v0.8.1 (Data S2; https://bitbucket.org/berkeleylab/checkv/src/master/).

### Identification of putative groundwater CPR phages

Three strategies were used to find out phages with the potential to infect groundwater CPR MAGs, including CRISPR spacers match, transfer RNA (tRNA) match, and nucleotide sequence homology search [18, 43, 44]. (i) *CRISPR spacers match*. The clustered regularly interspaced short palindromic repeat (CRISPR) was searched from 2790 CPR MAGs using metaCRT [45] (modified from CRT1.2) with default parameter. Extracted CRISPR spacers were further matched against groundwater viral contigs using BLASTn [46] (parameters: E value ≤ 1E-5, percentage identity of 95%, mismatches ≤ 1). (ii) *tRNA match*. The tRNA genes contained in groundwater viral contigs were identified by ARAGORN v1.2.38 [47] with the “−t” option, further aligned with the 2,790 CPR MAGs using BLASTn [46] (90% coverage and 90% identity). (iii) *Nucleotide sequence homology search*. Groundwater viral contigs were searched against 2,790 CPR MAGs based on shared genomic regions using BLASTn [46] (thresholds: 75% minimum coverage of the viral contig length, 70% minimum nucleotide identity, 50 minimum bit score, and 0.001 maximum e-value). Previous studies suggested this method yielded prophages [18]. The groundwater CPR phages were further clustered into CPR vOTUs using the same method mentioned above. All the host CPR MAGs were de-replicated at 95% average nucleotide identity (ANI) using dRep v3.2.2 [48] to generate estimated species-level host CPR OTUs (Operational Taxonomic Units; parameters: -comp 50 -con 10 -nc 0.30 -pa 0.9 -sa 0.95).

### Taxonomic assignment and lifestyle prediction of groundwater CPR phages

All the groundwater vOTUs (including those connected to host CPR MAGs) were further annotated using geNomad v1.7.3 [49] based on ICTV (International Committee on Taxonomy of Viruses) classification to get the taxonomic assignment. Referring to previous study [35], lifestyle of the groundwater CPR vOTUs was predicted using a strategy combined results of geNomad [49], CheckV [42], BACPHLIP [50] and nucleotide sequence homology search [18]. (i) For all the CPR vOTUs, integrated prophages identified by both geNomad and CheckV were considered as conservative temperate phages. (ii) For complete and high-quality CPR vOTUs, those with a probability above 90% in BACPHLIP predictions were considered as conservative temperate phages and virulent phages, respectively. (iii) Other CPR vOTUs that were connected to host CPR MAGs based on nucleotide sequence homology search were considered as potential temperate phages. (iv) Lifestyle of the remaining CPR vOTUs were regarded as unknown.

### Gene-sharing network analysis

The sequences of vOTUs recovered in this study, including the groundwater CPR vOTUs, were pooled to call open reading frames (ORFs) using Prodigal v2.6.3 [51] (parameters: -p meta -f gff -q -m). The resulting protein sequences were further input into vConTACT2 v2.086 [52] to run against the NCBI Prokaryotic Viral RefSeq v201 database (parameters: --rel-mode Diamond --pcs-mode MCL --vcs-mode ClusterONE). Phages were clustered based on the similarity of the shared protein clusters. The resulting network was visualized in Cytoscape v3.9.1 [53] (http://cytoscape.org).

### Abundance profiles

RPKM (Reads per kilobase per million mapped reads) were used to represent relative abundance of host CPR OTUs and their vOTUs. The RPKM of vOTUs were calculated using CoverM v0.6.1 (https://github.com/wwood/CoverM) to generate coverage profiles across samples (parameters: coverm contig --min-read-percent-identity 0.95, --min-read-aligned-percent 0.75, --contig-end-exclusion 0 and -m rpkm). Similarly, the RPKM of host CPR OTUs were also calculated using CoverM v0.6.1 to generate coverage profiles across samples (parameters: coverm genome --min-read-percent-identity 0.95, --min-read-aligned-percent 0.75, --contig-end-exclusion 0 and -m rpkm). Similar to VHR (Viral-to-Host Ratios), vOTUs/host CPR OTUs abundance ratios were calculated by dividing the RPKM of vOTUs by the RPKM of host CPR OTUs.

### Identification of auxiliary metabolic genes

All the 1162 groundwater CPR phages were rerun by VirSorter v2.1 (--prep-for-dramv) to generate the output files that DRAM-v required [43, 54]. Then the DRAM-v v1.3.5 [55] were used to recover putative AMGs. To be more conservative, only putative AMGs with auxiliary score < 4 were retained. Putative AMGs without gene descriptions were further annotated using GhostKOALA to get functional annotations (https://www.kegg.jp/ghostkoala/). In addition, putative AMGs associated with ribosome, peptidase, and proteasome, or still without gene descriptions, were discarded. Subsequently, a manual curation process was undertaken by checking the presence of viral hallmark genes or virus genes upstream and downstream of the selected AMGs.

## Results

### Viruses are widely distributed in groundwater ecosystems

To uncover viruses in groundwater ecosystems, a total of 82 metagenomic datasets derived from sequential filtration of groundwater in Northern California and Colorado [7, 8, 11] were recruited and analyzed (Data S1). Raw sequencing reads were trimmed and assembled into contigs, then VirSoter2 were used to successfully recover 39,741 viral contigs that were ≥ 5 kb in size. Subsequently, all the viral contigs were clustered into 22,160 viral Operational Taxonomic Units (vOTUs), representing approximately species-level taxonomy (Data S2). The genomic size of vOTUs varied from 5 to 624 kb, with most vOTUs having the genomic size from 5 to 25 kb (Fig. S1). According to the taxonomic results based on the latest ICTV classification, 87.3% of the vOTUs could be assigned taxonomy (Fig. S2). At class level, most (98.8%) of them were assigned to the class Caudoviricetes. Viruses in this class exhibit extremely high diversity and are widely distributed across various environments [17], with host ranges encompassing both bacteria and archaea. Additionally, the vast majority (97.1% and 97.5%, respectively) of vOTUs lacked taxonomic annotations at order- or family-level. Only 2.5% of vOTUs had taxonomic annotations at the family level, including Herelleviridae (0.83%), Mimiviridae (0.41%), Straboviridae (0.23%), Demerecviridae (0.13%), and Kyanoviridae (0.13%). Intriguingly, 167 vOTUs were assigned to the phylum Nucleocytoviricota, which was constituted of nucleocytoplasmic large DNA viruses (NCLDVs; also called giant viruses). NCLDVs possess complex genomes and virions that are similar in size to, or even larger than, small cellular organisms [56].

### The diversity and novelty of groundwater CPR phages

To explore the potential impacts of phages on CPR bacteria in groundwater ecosystems, the phage-CPR interactions were investigated. Previous studies had proposed that most CPR bacteria lack CRISPR [57], which plays important roles in phage defense, while they might have alternative strategies to resist phage infection [10]. Similarly, only 269 (9.64%) of the collected 2,790 groundwater CPR MAGs were predicted to possess CRISPR. Given this, three strategies (CRISPR spacers match, transfer RNA (tRNA) match, and nucleotide sequence homology search) were employed for phage-CPR interactions (see Materials and Methods for more details; Fig. S3), ultimately generating 14,120 pairs phage-CPR interactions (Fig. S4A; Data S3) and uncovering 1,162 phages capable of infecting 2,438 host CPR MAGs (Data S4; Data S5). The groundwater CPR phages were subsequently clustered into 644 CPR viral Operational Taxonomic Units (CPR vOTUs). Among the CPR vOTUs, the majority had the genomic size ranging from 5 to 30 kb (Fig. 1A). As for the taxonomic classification, 57.5% of the CPR vOTUs could be assigned taxonomic information (Fig. 1C). The vast majority (99.9%) of the CPR vOTUs with taxonomic information were assigned to the class Caudoviricetes, with 4.3% of them having family-level taxonomic annotations, including Herelleviridae (3.5%), Schitoviridae (0.3%), Straboviridae (0.3%), and Kyanoviridae (0.3%). In terms of lifestyle, it was quite interesting that 63.8% of the CPR vOTUs were considered as temperate phages (5.9% and 57.9% for conservative temperate and potential temperate, respectively), while only 0.8% of the CPR vOTUs were inferred as virulent phages (see Materials and Methods for more details; Fig. 1B). Such a large ratio of temperate-to-virulent suggested that groundwater host CPR MAGs were more prone to be infected by temperate phages, implying ingenious interactions between phages and host CPR MAGs in groundwater ecosystems.

**Figure 1.**
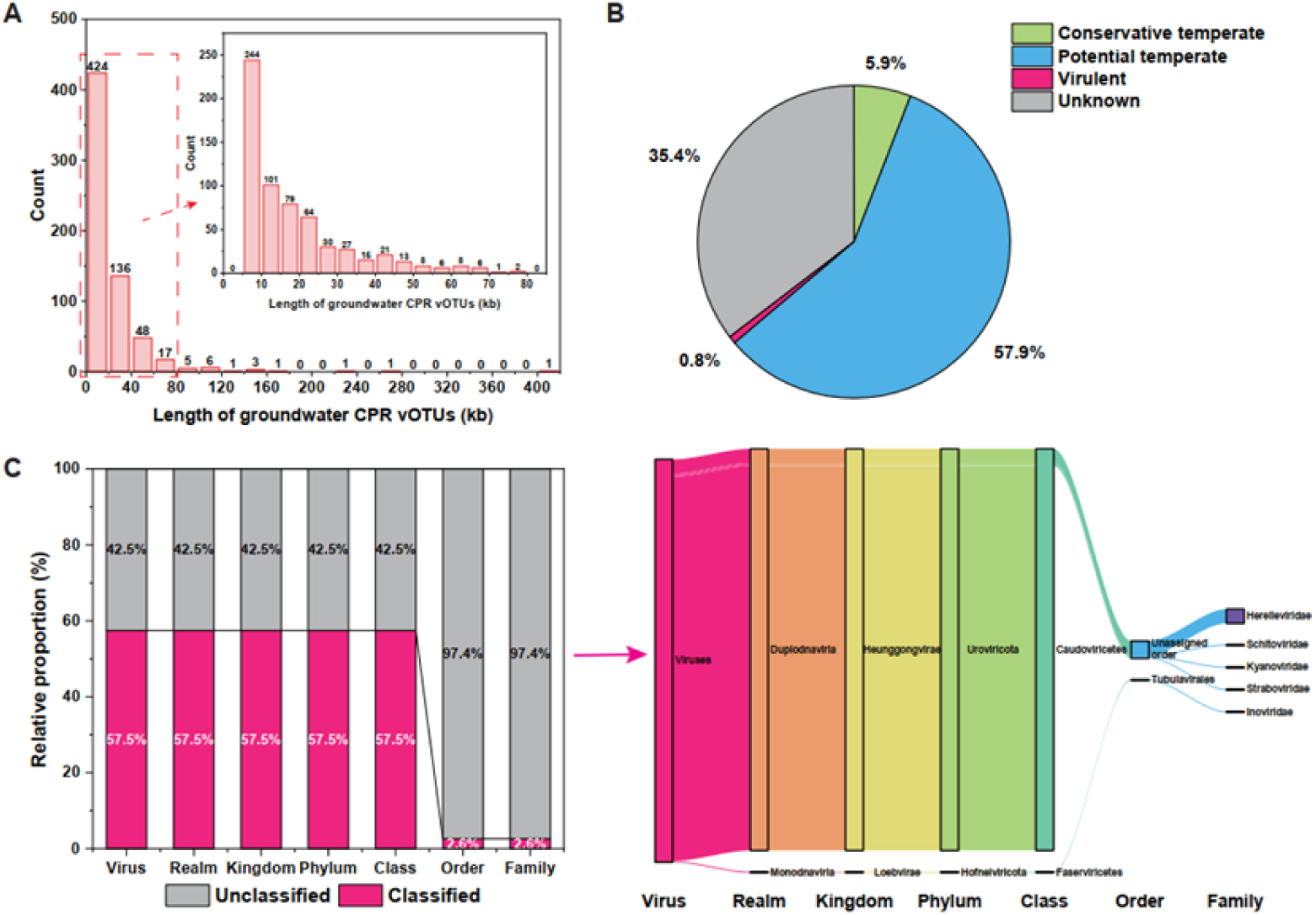
Overview of CPR vOTUs identified in groundwater ecosystems. (A) Length of groundwater CPR vOTUs. (B) Lifestyle proportion of groundwater CPR vOTUs. (C) Taxonomic affiliation of groundwater CPR vOTUs based on the latest ICTV classification. Bar plot shows the proportion of classified (pink) and unclassified (grey) vOTUs. Sankey plot shows taxonomic affiliation of classified CPR vOTUs.

To investigate the genomic similarity of groundwater CPR vOTUs, other vOTUs mined in groundwater ecosystems, and publicly available sequences, a gene-sharing network analysis was performed using vConTACT2 to produce genus-level viral clusters (VCs) (Fig. 2). In general, 391 (60.71%) CPR vOTUs, 11,804 (54.60%) groundwater other vOTUs, and 3,192 (91.15%) viral genomes from RefSeq v201, were grouped into 4,073 VCs. Among them, 32 VCs were exclusively comprised of groundwater CPR vOTUs, 3,300 VCs were solely formed by groundwater other vOTUs, and 533 VCs were purely composed of reference viral genomes. When assessing the genomic similarity between CPR vOTUs and other vOTUs mined in groundwater ecosystems, it was found that only 289 (44.88%) CPR vOTUs forming 178 VCs with other vOTUs, indicating a high degree of genomic novelty among CPR vOTUs. Surprisingly, only 2 (0.31%) CPR vOTUs could form 2 VCs with viral genomes from RefSeq v201, reflecting the huge and unexplored diversity of groundwater CPR vOTUs.

**Figure 2.**
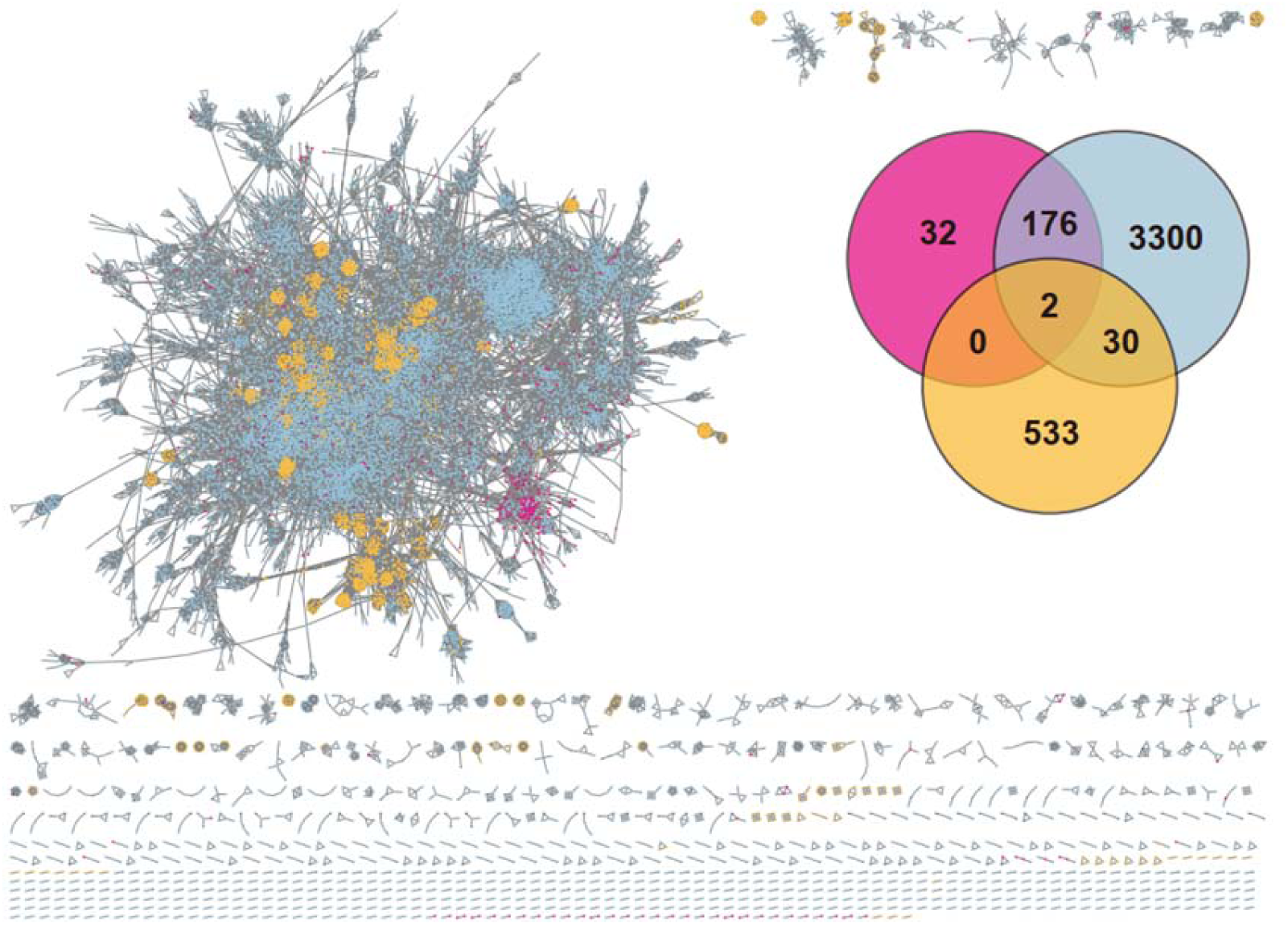
Relationship between vOTUs mined in this study and NCBI reference viral genomes based on gene-sharing network analysis. The nodes represent different viruses, while the shared edges indicate similarity based on shared protein clusters. Pink nodes represent groundwater CPR vOTUs, blue nodes represent other vOTUs discovered in this study, while the orange nodes represent NCBI reference viral genomes. Venn diagram shows the number of shared viral clusters among groundwater CPR vOTUs, groundwater other vOTUs, and NCBI reference viral genomes.

### Close interactions between host CPR MAGs and their phages in groundwater ecosystems

To better understand the complex interactions between phages and host CPR MAGs in groundwater ecosystems, both the capability of phages to infect host CPR MAGs and the susceptibility of host CPR MAGs to phage infection were examined. For the 1,162 CPR phages, 547 (47.1%) phages were exclusively connected to a unique host CPR MAG (termed specialist CPR phages), while 615 (52.9%) phages were connected to more than one host CPR MAGs (termed generalist CPR phages) (Fig. S4B). For 2,438 host CPR MAGs, only 310 (12.7%) host CPR MAGs were connected to a certain CPR phage (termed specialist host CPR MAGs), while 2,128 (87.3%) host CPR MAGs were connected to more than one CPR phage (termed generalist host CPR MAGs) (Fig. S4C). The host CPR MAGs were distributed across 14 classes, which were highly represented by the class Paceibacteria (n=1,178), Microgenomatia (n=656), ABY1 (n=297), Gracilibacteria (n=86) and WWE3 (n=84). With the exception of UBA1384 and CPR3, more than 75% CPR MAGs from other classes were predicted to be the host of phages (Fig. 3A). Phages connected to each class-level taxonomic affiliations of host CPR MAGs were dominated by temperate phages and generalist CPR phages (Fig. 3A).

**Figure 3.**
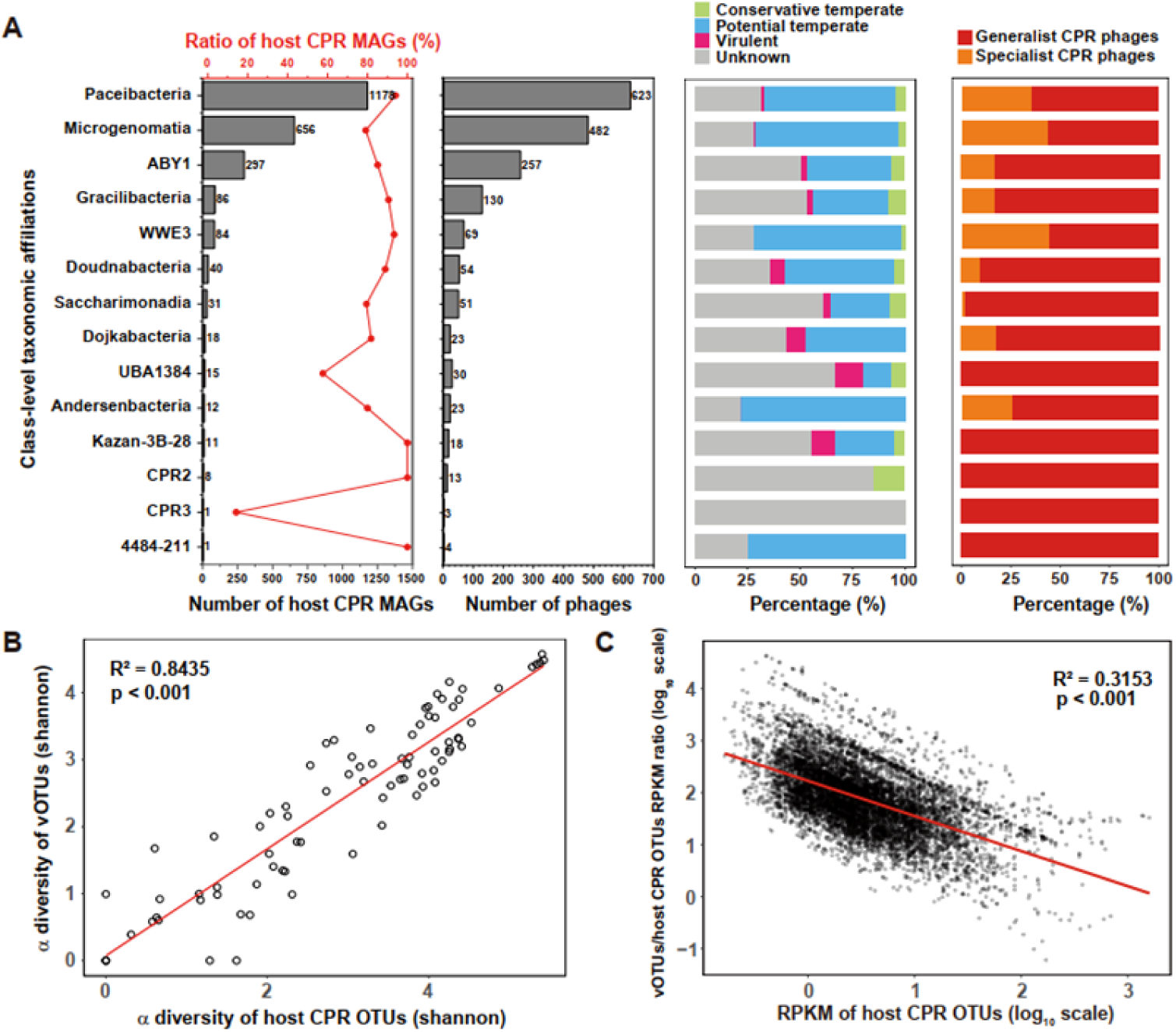
Interactions between host CPR MAGs and their phages in groundwater ecosystems. A, Number and ratio of host CPR MAGs, number of phages, lifestyle ratio of phages, and the ratio of generalist and specialist phages for each class-level taxonomic affiliations. B, Comparisons of α-diversity (Shannon index) between host CPR OTUs and their vOTUs. C, Correlation between the relative abundance of host CPR OTUs and vOTUs/host CPR OTUs abundance ratios.

Previous studies have shown that phages can regulate the structure and abundance of the microbial community by either microbial lysis (‘Kill the Winner’ hypothesis) or lysogenic infection (‘Piggybacking the Winner’ hypothesis) [58]. To further investigate the interactions between host CPR MAGs and their phages in groundwater ecosystems, we deduplicated these interactions to establish linkages between host CPR OTUs (Operational Taxonomic Units) and their vOTUs. Subsequently, a correlation analysis was performed on the profiles between host CPR OTUs and their vOTUs. There was a significantly positive correlation between the α-diversity (Shannon diversity) of host CPR OTUs and their vOTUs, indicating that their community structures were closely related in groundwater ecosystems (R^2^ = 0.8435, p < 0.001; Fig. 3B). Additionally, similar to VHR (Viral-to-Host Ratios), vOTUs/host CPR OTUs abundance ratios were significantly negatively correlated with the relative abundance of host CPR OTUs (R^2^ = 0.3153, p < 0.001; Fig. 3C). With the increase of the relative abundance of host CPR OTUs, vOTUs/host CPR OTUs abundance ratios decreased, referring to the ‘Piggyback-the-Winner’ (PtW) hypothesis. This hypothesis posits that more phages will choose to exploit their hosts by lysogenic infection rather than lytic infection at high microbial abundance [58]. Temperate phages can protect their hosts from being infected by others phages via superinfection exclusion [59] or alter the physiology of their host microbes via auxiliary metabolic genes [32], thereby enhancing the fitness and competitiveness of their host microbes. Therefore, when the abundance of host CPR OTUs in groundwater ecosystems is high, their vOTUs tend to enhance lysogenic infection to help them sustain the dominance in the microbial communities and maintain stable phage-CPR coexistence.

### Phages potentially promote symbiotic associations and metabolic exchange between host CPR MAGs and other free-living microbes

To further investigate how the groundwater CPR phages might affect host CPR MAGs, 355 phage-encoded putative auxiliary metabolic genes (AMGs) were predicted using DRAM-v (Data S6). Various indications suggest that CPR bacteria adopt a lifestyle that relies on other cells, either by deriving essential compounds from surroundings, or by episymbiotically attaching to larger host microbes to get nutrients [16].

Considering the requirement for attachment to other free-living microbes, it is unsurprising that CPR genomes typically encode numerous proteins associated with diverse cell-surface modifications, including pili, glycosyltransferase, and concanavalins (lectins) [16]. Interestingly, the groundwater CPR phages harbored five AMGs annotated as concanavalin A-like lectin/glucanases superfamily (Data S6). These proteins have been speculated to be involved in surface attachment of CPR bacteria to other free-living microbes [35]. What’s more, the groundwater CPR phages possessed 46 AMGs involved in O-Antigen nucleotide sugar biosynthesis, including genes encoding UDP-glucose 4-epimerase (*gal*E), dTDP-glucose 4,6-dehydratase (*rml*B), dTDP-4-dehydrorhamnose reductase (*rml*D), mannose-6-phosphate isomerase (*man*A), GDP-mannose 4,6-dehydratase (*gmd*) and GDP-4-dehydro-6-deoxy-D-mannose reductase (*rmd*) (Fig. 4; Data S6). The same as previous research [60], these AMGs encoded by phages may enrich rhamnose in the outer membrane lipopolysaccharide of host CPR MAGs by converting dTDP-4-oxo-6-deoxy-L-mannose to dTDP-L-rhamnose, and GDP-D-mannose to GDP-D-rhamnose, potentially contributing to surface adhesion and cell-cell aggregation of host CPR MAGs, thereby promoting symbiotic associations between host CPR MAGs and other free-living microbes.

**Figure 4.**
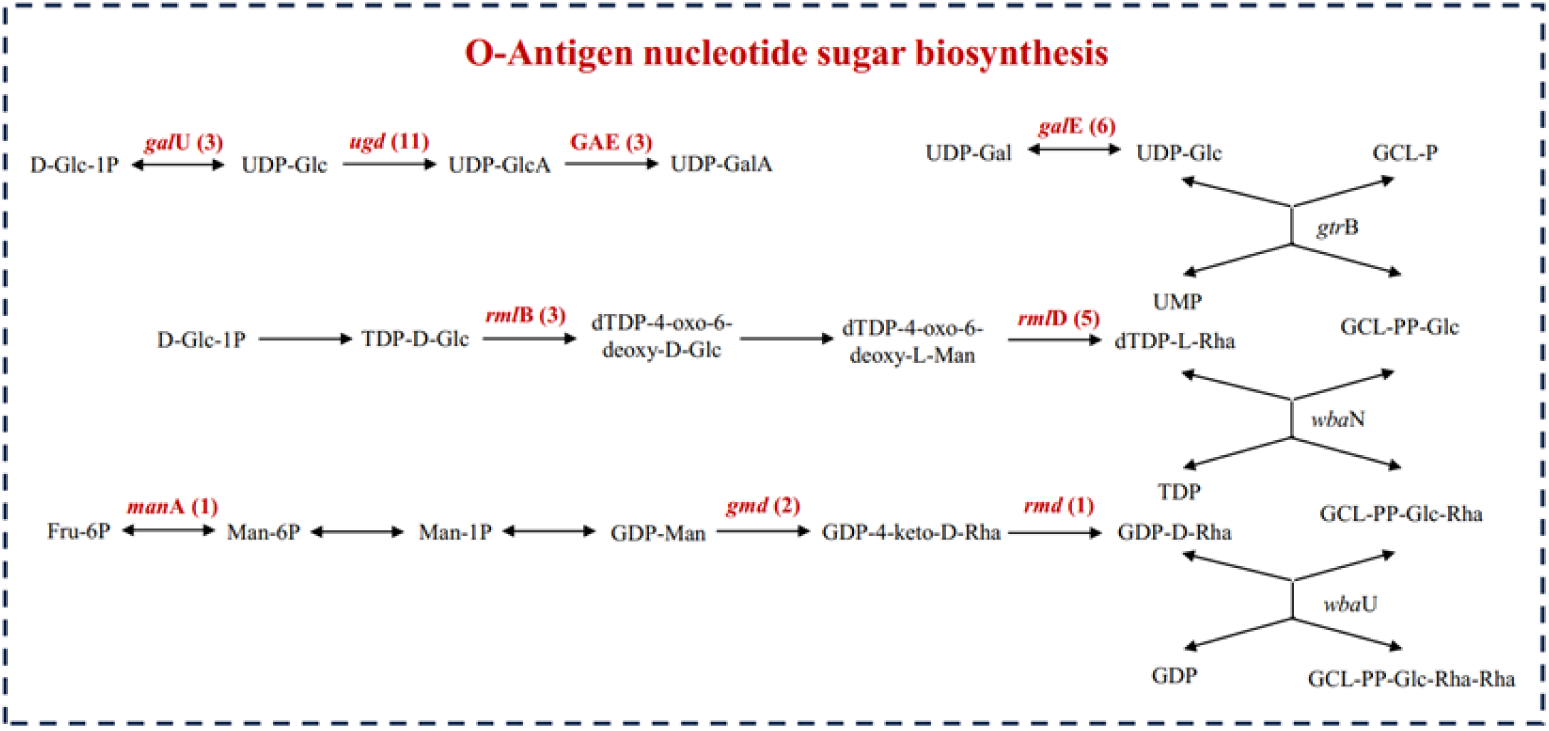
Putative auxiliary metabolic genes (AMGs) identified in groundwater CPR phages involved in O-Antigen nucleotide sugar biosynthesis. The phage-encoded AMGs are highlighted in red within these pathways, with Arabic numerals in parentheses indicating their respective quantities. GCL, glycosyl-carrier lipid.

Moreover, CPR genomes encode numerous transporters to fulfill the need for deriving essential compounds from surroundings [16]. It was ingeniously that the groundwater CPR phages encoded various AMGs associated with transport system, mainly including ABC transport system and P-type transporter (Table S1). ABC transport system utilizes the energy derived from ATP hydrolysis to actively transport nutrients (such as sugars, amino acids, inorganic ions) across concentration gradients into bacterial cells [61]. These AMGs encoding various transporters might also enhance the metabolic exchange between host CPR MAGs and other free-living microbes.

Besides, the groundwater CPR phages also encoding P-type transporters which can utilize the energy generated by ATP hydrolysis to transport specific ions (such as Ca^2+^, Mg^2+^, Cu^2+^, Cu^+^, etc.) across membranes [62], making them crucial for regulating ion concentration and osmotic pressure across the cell membrane. These ions also serve as essential cofactors for certain enzymes involved in bacterial growth and metabolism [63]. In oligotrophic groundwater ecosystems, through AMGs associated with transporters, phages might not only enhance the uptake of essential nutrients from surroundings, but also maintain ionic homeostasis across the cell membrane, thereby enhancing the environmental adaptability and survival ability of host CPR MAGs.

### Phages potentially expand metabolic capabilities of host CPR MAGs

Moreover, the groundwater CPR phages tended to encode AMGs associated with metabolism, mainly involved in Carbohydrate metabolism (n=113), Animo acid metabolism (n=85), Metabolism of cofactors and vitamins (n=68), Glycan biosynthesis and metabolism (n=56), Energy metabolism (n=55), and Nucleotide metabolism (n=43) based on KEGG database (Fig. S5).

For carbohydrate metabolism, five AMGs were annotated as concanavalin A-like lectin/glucanases superfamily which was associated with phage-encoded glycoside hydrolase, potentially involving in pectin cleavage [26]. The carbon sources for CPR bacteria typically consist of complex carbon compounds derived from degraded plant or microbial biomass [16]. Therefore, the ability to degrade these complex compounds into simpler monomeric forms is essential for them. By cleaving polymers into monomers, these glycoside hydrolases may facilitate carbon utilization of host CPR MAGs in oligotrophic groundwater ecosystems. What’s more, the groundwater CPR phages encoded various AMGs involved in glycolysis, pentose phosphate pathway and pyruvate metabolism (Fig. 5). Most of CPR bacteria lack complete glycolysis and pentose phosphate pathway, usually compensating for the impaired central carbon metabolism through metabolic shunt [16]. A previous study suggested that viruses could compensate for the incomplete central carbon metabolism (mainly pentose phosphate pathway) of DPANN archaea through AMGs [33]. This inspired our hypothesis that phages might significantly enhance the central carbon metabolism capacity of host CPR MAGs via AMGs, reducing their dependence on metabolic shunt. Certain CPR bacteria are abundant in acetate-amended groundwater, implying their ability to utilize acetate [7, 8]. Phage-carried AMGs encoding acetate kinase and aldehyde dehydrogenase might reversely promote acetate utilization, potentially explaining the proliferation of host CPR MAGs following acetate stimulation.

**Figure 5.**
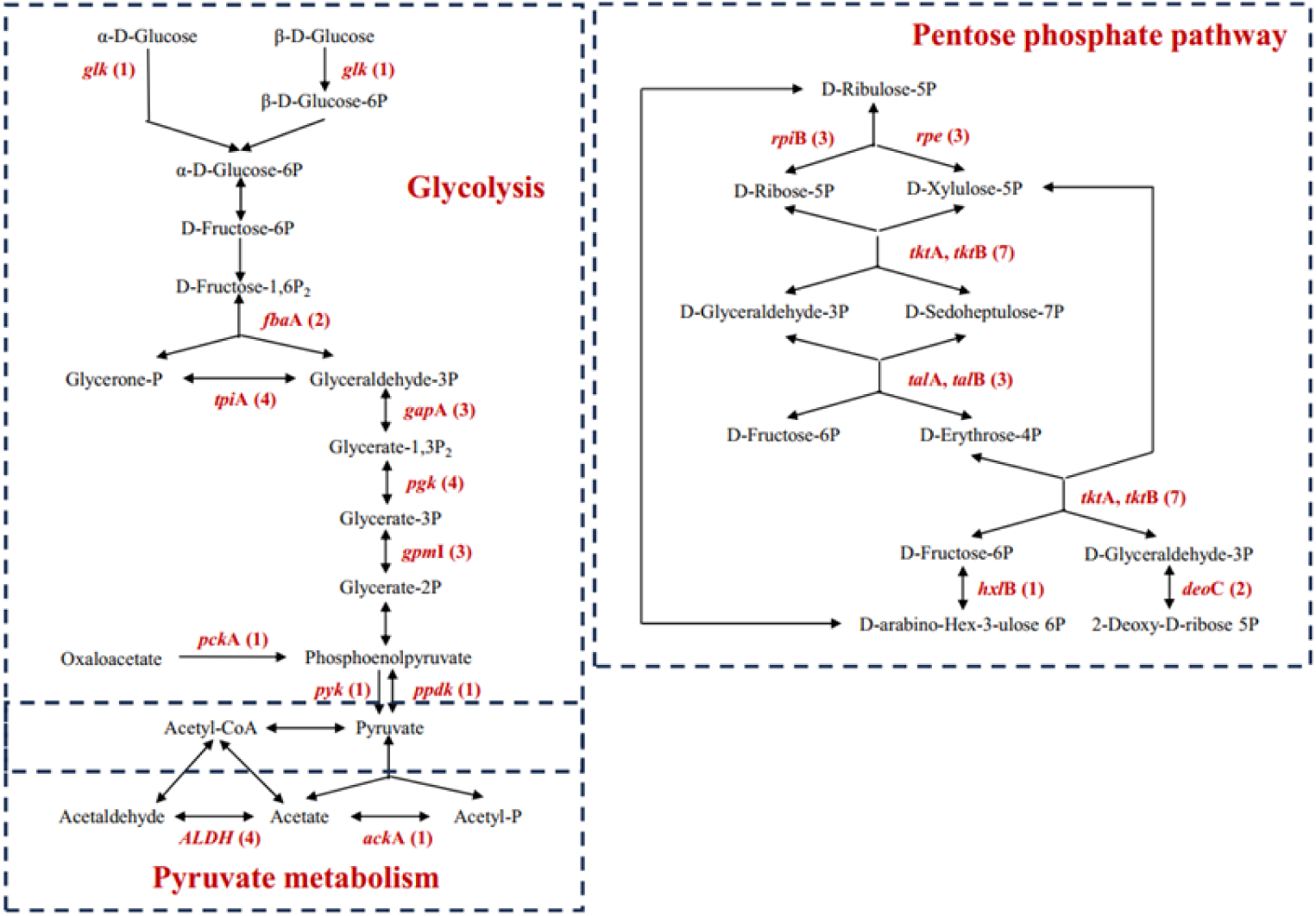
Putative auxiliary metabolic genes (AMGs) identified in groundwater CPR phages involved in glycolysis, pentose phosphate pathway and pyruvate metabolism. The phage-encoded AMGs are highlighted in red within these pathways, with Arabic numerals in parentheses indicating their respective quantities.

For metabolism of cofactors and vitamins, the groundwater CPR phages encoded various AMGs involved in nicotinate and nicotinamide metabolism, folate biosynthesis, and one carbon pool by folate (Fig. S6). Nicotinate and nicotinamide are precursors of NAD, which act as electron carriers in glycolysis, tricarboxylic acid cycle, and oxidative phosphorylation, participating in energy generation and conversion [64]. Folate is an important cofactor that mediates the transfer of one carbon unit [65]. It participates in the de novo synthesis of purines and pyrimidines, as well as the synthesis of various amino acids (glycine, serine, methionine, etc.), making it crucial for the growth and proliferation of bacterial cells. There were also two AMGs identified as cysteine desulfurase which could decompose L-cysteine to L-alanine and sulfane sulfur [66], thus potentially involving in the biosynthesis of a variety of cofactors (including Fe-S clusters in proteins, thionucleosides in tRNA, thiamine, biotin, lipoic acid, molybdopterin, and NAD) (Data S6). Many enzymes encoded in CPR bacteria require cofactors, yet pathways for their biosynthesis were rarely identified [16]. In this context, phage-encoded AMGs might partially alleviate the challenges host CPR MAGs face in cofactors and vitamins metabolism.

For energy metabolism, there were five AMGs involved in oxidative phosphorylation, encoding F-type H^+^/Na^+^-transporting ATPase subunit alpha, beta, and gamma (Data S6). F-type H^+^/Na^+^-transporting ATPase utilizes transmembrane proton gradient to drive the production of ATP, which serves as the primary energy source for various biological processes in bacterial cells [67]. Additionally, the groundwater CPR phages encoded numerous AMGs involved in animo acid metabolism and nucleotide metabolism (Fig. S7; Data S6). CPR bacteria appear to have defects in the biosynthesis of amino acids and nucleotides, often needing to salvage them from external sources [16]. However, phage-encoded AMGs might alleviate these deficiencies and reduce their dependence on surroundings.

## Discussion

In this study, we mined phages capable of infecting host CPR MAGs from 82 groundwater metagenomic datasets. The taxonomic assignment and gene-sharing network analysis indicated rich diversity and great novelty of the groundwater CPR vOTUs. In groundwater ecosystems, host CPR OTUs and their vOTUs formed intricate interactions, conforming to ‘Piggyback-the-Winner’ (PtW) hypothesis. We also identified diverse AMGs in CPR phages which might facilitate the symbiotic associations and metabolic exchange between host CPR MAGs and other free-living microbes and expand metabolic capabilities of host CPR MAGs.

Viral lifestyle strategies are central to the ecology of virus-host interactions, mainly through the lytic and lysogenic replication cycles, referring to ‘Kill the Winner’ (KtW) hypothesis and ‘Piggyback-the-Winner’ (PtW) hypothesis, respectively [58]. Lytic infection leads to the reproduction of viral progeny, the lysis of host cells, and the release of the cellular dissolved organic matter into the surrounding environment [31]. The corresponding KtW hypothesis suggests that higher host microbial abundance is associated with stronger lytic infection, leading to higher VHR [32]. In this case, viruses increase the availability of resources by persistently killing the dominate microbes, promoting the growth of relatively disadvantaged species, thereby enhancing the diversity of the microbial community [58]. Lysogenic infection is chosen by temperate viruses to integrate into, and replicate with, the genome of host microbes by forming a mutualistic relationship [68]. The corresponding PtW hypothesis suggests that higher host microbial abundance enhances lysogenic infection and reduced VHR [59]. In this context, viruses confer competitive advantages to the dominant microbes through superinfection exclusion or auxiliary metabolic genes, inhibiting the growth of relatively disadvantaged species, thereby reducing the diversity of the microbial community [58]. We found that host CPR MAGs were predominantly infected by temperate phages in groundwater ecosystems, and the abundance profiles between host CPR OTUs and vOTUs conformed to the PtW hypothesis. Given that groundwater ecosystems are typical oligotrophic habitats, and CPR bacteria had high relative abundance in the collected groundwater samples, these findings are consistent with the previously reported view that lysogenic infection was generally considered to be more prevalent and important in harsh environments [32, 58]. From the perspective of phages, the lack of CRISPR in CPR bacteria provides opportunities and convenience for their lysogenic infection. From the perspective of CPR bacteria, they face a challenge of relying on external resources for survival due to deficiencies in core metabolic capabilities. Therefore, they need to enhance lysogenic infection to maximize their competitive advantages, thus enabling them to maintain the dominance in the microbial communities. Overall, lysogenic infection is a strategy co-evolved by CPR bacteria and their phages in groundwater ecosystems.

Groundwater ecosystems are typical oligotrophic habitats with nutrients in low concentration and limited diversity [69], driving CPR bacteria residing in these habitats to streamline their metabolic capabilities and biosynthetic pathways. A previous study suggested that most of the genomes of CPR bacteria lacked complete glycolysis and pentose phosphate pathway, compensating for the impaired central carbon metabolism through metabolic shunt [16]. Moreover, due to the lack of complete biosynthetic pathways for cofactors, amino acids, and nucleotides, many CPR bacteria need to acquire these nutrients from external sources [16]. Intriguingly, we identified various AMGs in CPR phages which might play interesting roles in the interactions between host CPR MAGs and their phages in groundwater ecosystems (Fig. 6). As symbionts that rely on other free-living microbes for essential nutrients, CPR bacteria can attach to other larger microbes though various cell surface modifications [16]. The groundwater CPR phages encoded numerous AMGs associated with concanavalin A-like lectin/glucanases superfamily and O-Antigen nucleotide sugar biosynthesis, which could enhance surface adhesion of host CPR MAGs. These AMGs, along with AMGs associated with ABC transport system and P-type transporter, might strengthen episymbiotic associations and metabolic exchange between host CPR MAGs and other free-living microbes. This would, in return, increase the likelihood of CPR phages encountering potential host microbes, thus enhancing their infectivity. The groundwater CPR phages also harbored numerous AMGs, which not only alleviate metabolic deficiencies in host CPR MAGs but also enhance their uptake of essential nutrients from surroundings. With the assist of these AMGs, the growth, proliferation, and survival ability of host CPR MAGs would be promoted, thereby maintaining stable phage-CPR coexistence.

**Figure 6.**
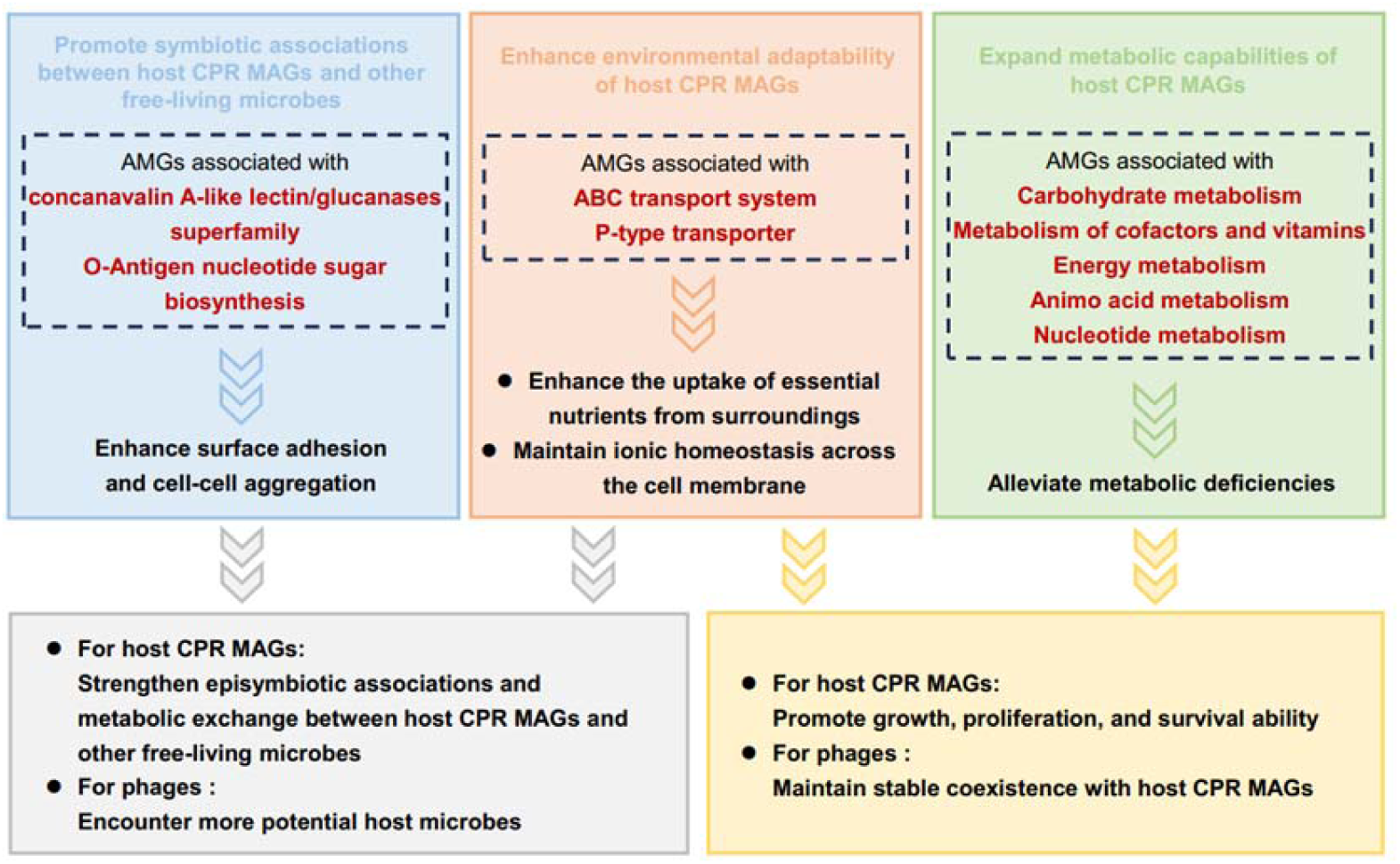
Schematics of the impact of auxiliary metabolic genes (AMGs) on the interactions between host CPR MAGs and their phages in groundwater ecosystems.

In conclusion, our study revealed the diversity and novelty of phages with the potential to infect host CPR MAGs in groundwater ecosystems. We elucidated the complex interactions between host CPR MAGs and their phages, and found that phages potentially promoted the symbiotic lifestyle and metabolic potential of host CPR MAGs through AMGs. These results provide valuable insights into the ecological roles of phage-CPR interactions and their co-evolutionary strategies in groundwater ecosystems. While further laboratorial and in situ experiments are required to validate these findings.

## Author contributions

## Supplementary material

## Data availability

